# Energy-harnessing problem solving of primordial life: modeling the emergence of catalytic host-parasite life cycles

**DOI:** 10.1101/2021.02.03.429491

**Authors:** Bernard Conrad, Magnus Pirovino

**Affiliations:** Genesupport, Avenue de Sévelin 18, CH-1004 Lausanne; Opiro, Landstrasse 40, FL-9495 Triesen

## Abstract

All life forms on earth ultimately descended from a primordial population dubbed the last universal common ancestor or LUCA via Darwinian evolution. Extant living systems share two salient functional features, a metabolism extracting and transforming energy required for survival, and an evolvable, informational polymer – the genome – conferring heredity. Genome replication invariably generates essential and ubiquitous genetic parasites. Here we model the energetic, replicative conditions of LUCA-like organisms and their parasites, as well as adaptive problem solving of host-parasite pairs. We show using the Lotka-Volterra equations that three host-parasite pairs – individually a unit of a host and a parasite that is itself parasitized – are sufficient for robust and stable homeostasis, forming a life cycle. This catalytic life cycle efficiently captures, channels and transforms energy, enabling dynamic host survival and adaptation. We propose a Malthusian fitness model for an original quasispecies evolving through a host-parasite life cycle.

## Introduction

One common definition of life posits that it represents a self-sustained chemical system capable of evolution (*1*). While there is no general consensus on what life is and how it originated (*2*), it is widely appreciated that all life forms on earth ultimately descended from a theorized, primordial population named LUCA via Darwinian evolution (*3, 4*). Extant living systems share two key functional components, a metabolism extracting and transforming energy required for survival, and an evolvable, informational polymer conferring heredity, i.e. the genome. Two competing – although not mutually exclusive – concepts stipulate that either primitive self-replicators (genetics-first) or self-reproducing and evolving proto-metabolic networks (metabolism-first) were critical at the origin of life (*5, 6*).

Genetic parasites are an integral feature of genome replication, and constitute the most abundant and diverse biological entity on earth (*7*). Arguably the emergence and persistence of parasites is unavoidable, since parasite-free states are evolutionary unstable, and microbial populations cannot clear parasites yet escape extinction through Muller’s ratchet (*8, 9*). They parasitize all cellular life forms including LUCA, with the possible exception of certain obligate intracellular bacteria with a reduced genome leading a parasitic life style (*10*). Attempts at reconstructing the LUCA parasitome revealed a remarkably large virome comprising the extant viruses of bacteria and archaea (*11*). Intriguingly, ancient bacterial symbionts undergoing massive genome erosion repeatedly experienced extinction and replacement by pathogenic microbes (*12*). Collectively, this suggests that evolution of life essentially is host-parasite co-evolution (*7*). Indeed, host-parasite co-evolution is considered among the dominant drivers of biological diversity over the last 3.5 billion years (*13*). It further implies a life cycle of host-parasite interactions that represent a temporal trajectory of competition, cooperation, and replacement by fresh parasites once the ancient symbiont genome is eroded (*13, 14*), (Methods; suppl. Figs. 1 and 2). As a corollary, a host exposed to a given parasite encounters a parasite that is it-self parasitized, forming what we henceforth call a host-parasite unit (Methods).

A host therefore is a dynamic host-parasite unit with multiple, changing partners that evolves along a temporal axis implying competition and cooperation. We set out to implement this paradigm for a LUCA-like founder population using the Lotka-Volterra equations. Lotka (*15*) and Volterra (*16*) independently modelled prey-predator interactions, both concluding that populations would oscillate because of this interaction. The respective equations became known as Lotka-Volterra equations, which reduced the then popular thinking of complex food chains to the interaction of only two species controlling each other in a cyclic manner (*17*).

## Results

In what follows we will initially formalize mathematical proofs for host-parasite interactions with the help of the Lotka-Volterra equations. We then go on to address virulence – as a proxy for an actively reproducing parasite – in relation to the energy made available to a host at a given moment, since energy is at the core of the metabolic component of living systems.

Assuming a LUCA or pre-LUCA host-parasite unit composed of a host infected by bacteria (a heterogeneous, prokaryotic population with a genomic complexity similar to extant bacteria and archaea) that are parasitized by a phage population (LUCA or pre-LUCA virome), and if this host serves as a stable habitat nourishing the bacterial population *X*, the phage population *Y* stabilizes *X* in its host, if *X* is a necessary nutrient for *Y*. Two lines of proof will be developed, firstly for the stabilization of the interacting prokaryote-phage population, and secondly to demonstrate that the fitness of the host is synchronized with that of its prokaryote-virus pair.

The proof outline for stabilization of the prokaryote-phage populations is as follows; populations *X* and *Y* fulfil the classical predator-prey relationship as described by the Lotka-Volterra equations 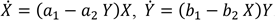. Based on the Lotka-Volterra calculus we know that populations *X* and *Y* fluctuate around their equilibria 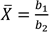 and 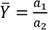. Both equilibria, 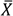 and 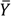, are independent of their initial population size as long as they start with positive values. The association of a dedicated phage with a given prokaryote is a very effective and robust taming strategy for a parasitized host. If there is no phage (i.e. *Y* = 0), then we conclude in accordance with the above Lotka-Volterra equations 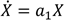, with *a*_1_ > 0. Consequently, the growth of prokaryote *X* is unlimited, as long as the host habitat nourishes it. However, if the host loses the ability to promote this unlimited growth, it will be destroyed during the process. If virulent and active, a bacterial parasite without phage potentially always exerts deleterious effects on the host.

The proof outline for fitness synchronization between the host and the prokaryote-virus pair is as follows; from the Lotka-Volterra calculus it follows that the stabilized equilibria populations of prokaryote 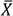 and their phages 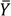 are proportional to the size of their habitat (i.e. to the number of possible encounters between members of *X* and *Y*). As a corollary, the size of their habitat itself is proportional to the number of hosts. Thus, host population and equilibria population sizes of prokaryotes and their viruses are interlinked, implying that host fitness can be synchronized with the fitness of the equilibria populations 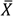 and 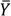. In contrast to other quantitative models, the Lotka-Volterra equations used here do not include all replicators involved – i.e. the host, the prokaryotes and their viruses – but only model the replication of prokaryotes (including viruses). As discussed, the growth rates of parasites (plus viruses) and hosts are synchronized implicitly, via the growth rates of the parasites’ equilibria populations. While this confirms previous findings showing that parasites stabilize their host (*8, 9*), the trio of a host interacting with a parasitized parasite expands the probability space of encounters – the host is modelled intrinsically – reaching far beyond previous models.

Living systems are thermodynamic entities (*18*), sometimes referred to as energy harnessing device making copies of itself (*19*). Boltzmann pointed out that “the fundamental object of contention in the life-struggle, in the evolution of the organic world, is available energy”, and Lotka stated that “in the struggle for existence, the advantage must go to those organisms whose energy-capturing devices are most efficient in directing available energy into channels favorable to the preservation of the species” (*20*).

For these reasons, we modelled virulence in relation to energy (Methods). In what follows we will show in a gradual process that two energies and three trios of host-parasite units are required to attain robust homeostasis (the host is modelled intrinsically, trios may represent temporally successive states of the same unit; suppl. Fig. 2). At least two energy forms counter the natural fitness decay related to mutation, genetic drift (*21*) and environmental changes; those energies are also required for reproduction. Both can be viewed as proxies for the Malthusian fitness of the quasispecies. We based our simulation on a simple energy-virulence relation (Fig. 1). Both energies, *E* = *E*_1_ and *E* = *E*_2_, have two levels through which they regulate the virulence of parasites sensitive to them. If *E*_1_ is abundant 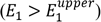, the virulence of (*E*_1_– sensitive) parasites is zero: *v* = 0. If *E*_1_declines, the parasite remains dormant (*v* = 0), until energy drops below a lower level 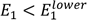. This allows the parasite to be reactivated and to acquire maximal virulence (*v* = 1). If in a second time point the host’s energy level rises again, the parasite remains virulent until *E*_1_ reaches the upper level again, which is the threshold allowing the host to tame all of its *E*_1_-sensitive parasites (*v* = 0), and so on.

**Fig. 1.**
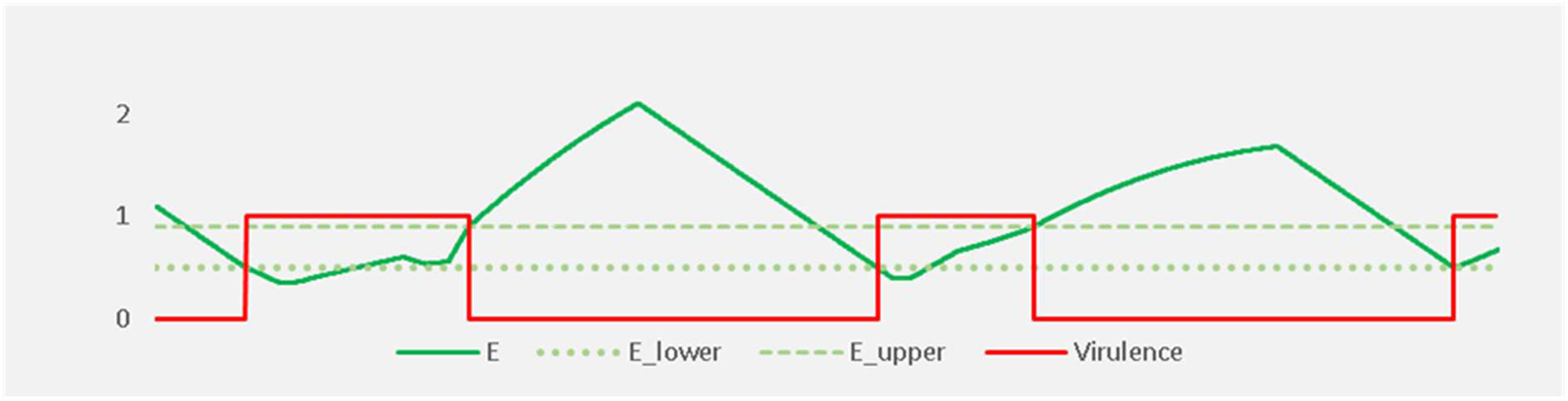
Host-parasite energy-virulence model. It illustrates the relationship between the energy level *E* = *E*_1_ of the host and virulence of its parasites. Both energies, *E* = *E*_1_ and *E* = *E*_2_, have two levels, *E*^*lower*^ and *E*^*upper*^, through which they «regulate» the virulence of parasites that are sensitive to them.

In Fig. 2 we extend this model by adding a parasite (viruses e.g. bacteriophages) to the parasite (prokaryote e.g. bacteria or archaea) to form the host-parasite unit, and limit it to one single energy (*E* = *E*_1_). This yields the “one energy, one host-parasite homeostasis model” (for the mathematical formalism and description see Methods). This simple model leads to an initial level of homeostasis – in the host or more generally in the whole quasispecies – as long as enough new operating power *E* can be catalyzed by some symbiotic host-parasite unit (prokaryote *X* and virus *Y*), and whenever the system skips into a problematic zone.

**Fig. 2.**
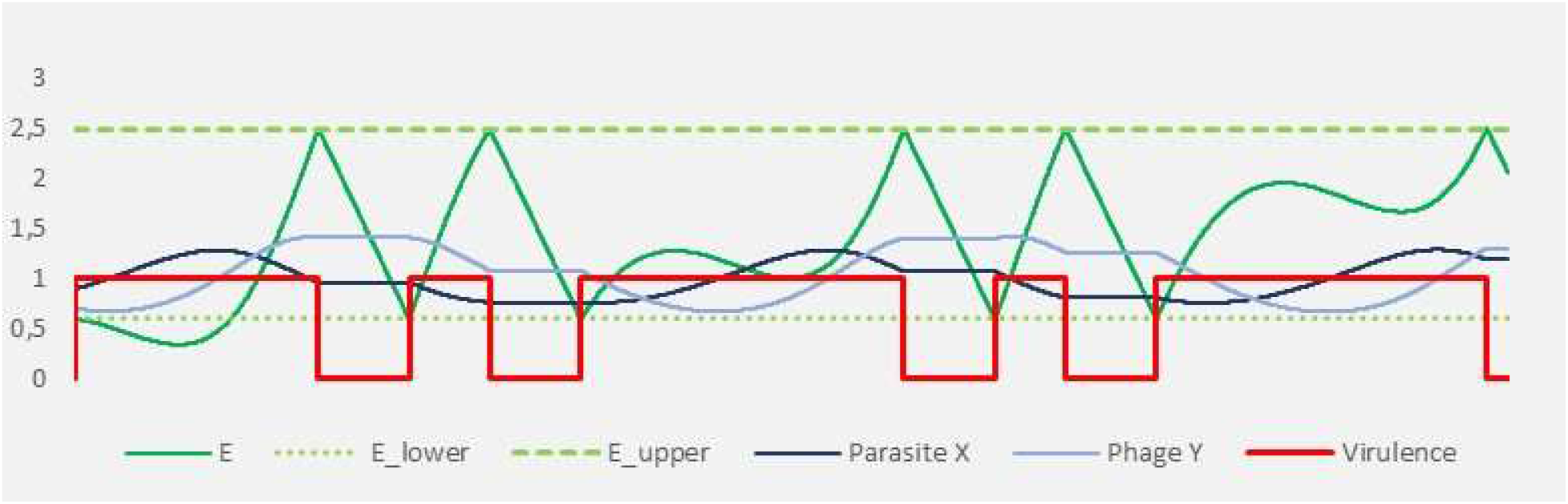
One energy, one host-parasite unit homeostasis model. The model is extended by adding a parasite (phage *Y*) to the parasite *X* (bacteria) to form the host-parasite unit, limited to one single energy (*E* = *E*_1_) that has two levels, *E*^*lower*^ and *E*^*upper*^.

This elementary homeostasis model – based on one single energy *E* = *E*_1_ regulating virulence and one host-parasite pair – still is considerably unstable. In case the symbiotic parasite-phage pair lacks its ability to catalyze new energy *E*, for instance because of fitness erosion or an environmental change, the system loses its problem fixing ability and breaks down. In its problematic zone (*E* < *E*^*lower*^), the system is in search mode, i.e. in the resolution search process (RSP; evolutionary problem solving discussed in Methods), without obvious problem fixing capacity. A new symbiotic prokaryote-virus pair capable of catalyzing new operating power *E* has to be selected in the parasite pool from the habitat’s available components. In this problematic zone, the system already has a low level of operating power *E* that is further decreasing, as long as no new solution is found. Accordingly, the system literally runs out of fuel until a new “fuel generator” can be identified. The inherently slow RSP needs to buy time in order to find a new, rapid problem fixing process (PFP; see Methods). In conclusion, the system requires a second fallback energy system that is not available in the one energy model.

Furthermore, in order to attain stable homeostasis using two energy levels (*E*_1_, *E*_2_), a quasispecies necessitates to codify the relevant transformations of these energy forms into one another. This is accomplished by combining three individual host-parasite units (“two energies, three host-parasite units homeostasis model” in Figs. 3a and 3b, in two different time scales; for mathematical derivation and description see Methods). Energy transformation by host-parasite units is supported by a vast body of literature, to name a few examples, i) the origin of the CRISPR-cas adaptive immune system in bacteria and archaea as a result of encounters with foreign, parasitic nucleic acids (*22*); ii) the acquisition of adaptive immune V(D)J recombination in jawed vertebrates via parasitic *ProtoRAG* transposable elements (*23*); iii) the emergence of 5-methylcytosine (5mC) in many species driven by the requirement to silence parasitic elements (*24*); iv) programmed cell death (PCD) likely originating from antiparasitic defense mechanisms activated when immunity fails (*25*); v) DNA parasites that led to the rapid, extensive intron gain during eukaryotic evolution (*26*), and finally, vi), the emergence of synaptic memory via Arc, a master regulator of synaptic plasticity that is related to a retrotransposon gene (*27*).

**Fig. 3.**
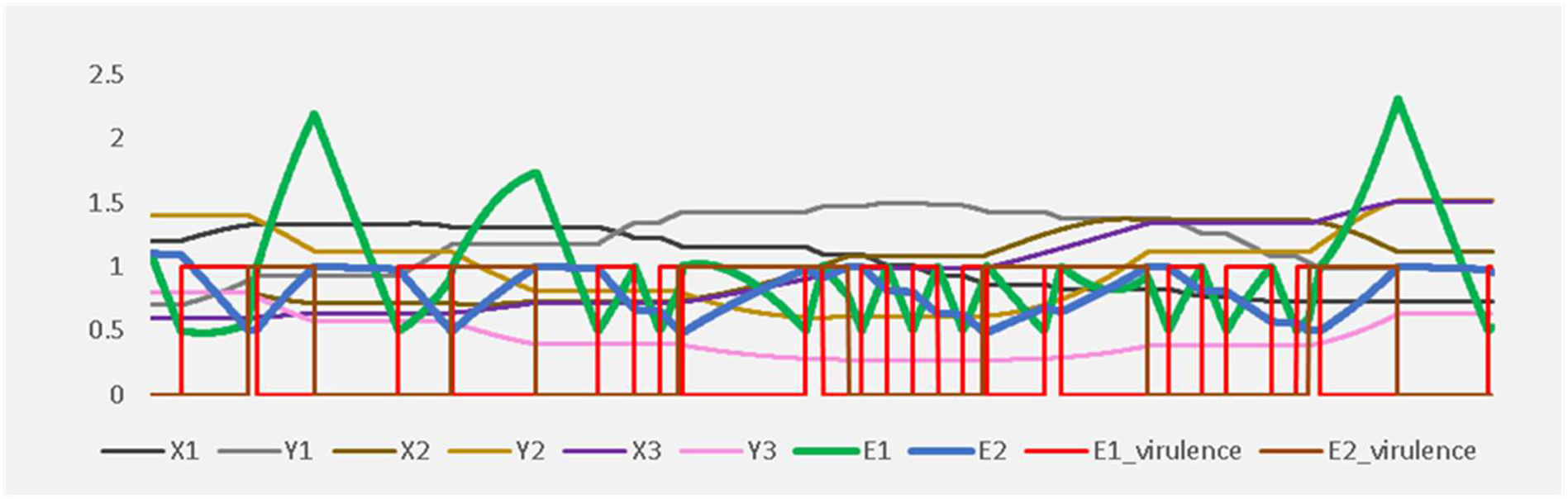
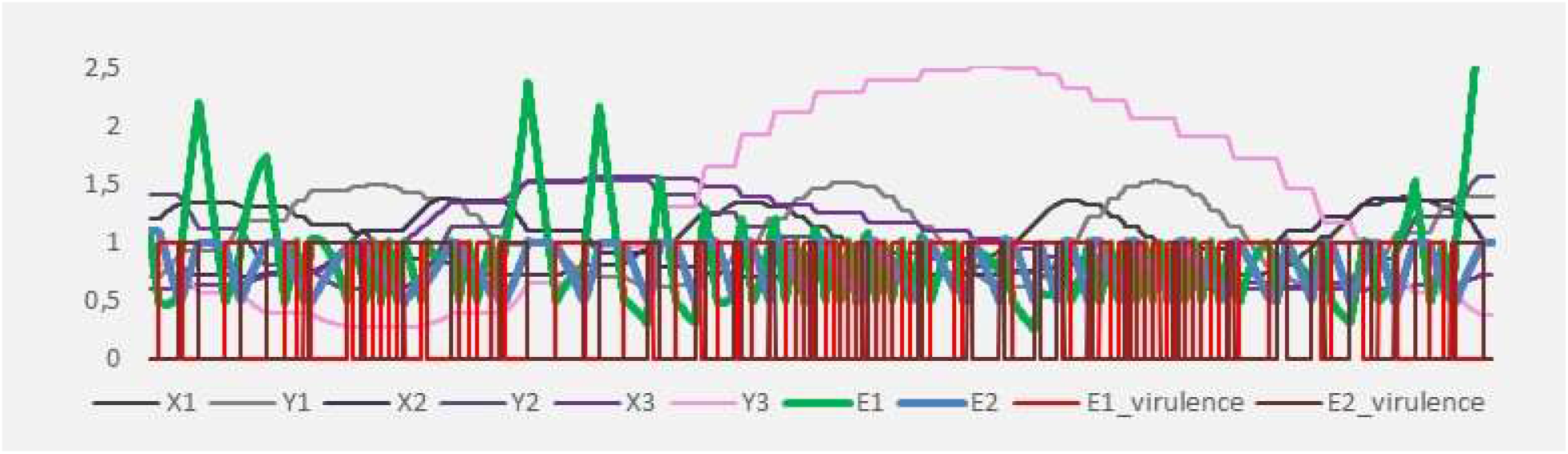
Two energies, three host-parasite units homeostasis model. Fig. 3A with timescale A, and Fig. 3B with timescale B. The model is extended by adding three host-parasite units (*X*_1-3_, *Y*_1-3_), and two energy levels (*E* = *E*_1_ and *E* = *E*_2_). The virulence of (*X*_1_, *Y*_1_) is regulated by the operating energy *E*_1_. (*X*_2_, *Y*_2_) and (*X*_3_, *Y*_3_) are *E*_2_– sensitive.

In conclusion, the main advantage of the “two energy homeostasis” over the “one energy homeostasis” lies in its robustness against environmental fluctuations and against functional defects of the symbiotic host-parasite units. If the system enters into a problematic zone, the slow RSP now has more time to find a new PFP, since this process can rely on a second fallback energy.

A more in-depth consideration of the individual nature of the energy transformation steps is warranted, since energy transformation is a salient feature of the model. Notably, i) Process 2 associated with transforming *E*_1_ → *E*_1_, is exemplified by the innate immune system (*28*); ii) 5mC-methylation is a paradigm for Process 3 and *E*_1_ → *E*_2_ transformation (*24*); finally, iii) Process 1, implying *E*_2_ → *E*_1_, is substantiated by acquired immunity and PCD (*22, 23, 25*), and by parasite-mediated cis-regulatory transcriptional regulation (*29*). In support of an energy transformation paradigm, evidence for forth and back shuttling of parasite components between parasites and the host defense systems has recently been provided (*30*). Moreover, the fact that fresh parasites are repeatedly recruited once ancient symbionts degenerated (*12*), a fate that appears to be the general rule for obligate endosymbionts (*14*), lends substantial credit to the idea of a host-parasite life cycle. We furthermore show that this host-parasite life cycle efficiently recycles energy from the fundamental process of replication-inherent parasites, whereby it captures, channels and transforms energy that can be used for further evolutionary purposes.

How can this life cycle of three host-parasite units then be integrated into a broader evolutionary genetic context? The core metrics of evolutionary genetics is fitness (*31, 32*). We propose a model in ten steps centered on fitness management of a quasispecies. Step 1: a quasispecies faces an evolutionary problem (i.e. mutation, genetic drift, environmental change) when the range of Malthusian fitness values for all its members is negative (see Methods). The problem resolution unfailingly implies an increase in fitness variance (*var*(*m*)). Step 2: a reduced fitness potentiates the environmental exposure, further decreases the mean fitness but increases the fitness variance. The increase in fitness variance in turn kick-starts the RSP, providing a potential solution. Step 3 specifies the counter forces mobilized by an eroded fitness, namely the two energy forms *E*_1_ and *E*_2_. Step 4 describes the fact that, in order to increase fitness, a quasispecies can only codify and enhance physical effects which are already in place («catalytic principle»). Step 5: based on step 4 the quasispecies has merely problem solving solutions at hand that are inherent to its own replication system, namely parasites (see «catalytic principle»). Those parasites subsequently codify and amplify the RSP in step 5. Parasites by their very nature gain in virulence whenever the host fitness is eroded, and parasite amplification enhances the process of fitness disintegration and lastly incrementally boosts fitness variance. In step 6, this RSP amplification leads to recruitment of those elements among the parasitic waves that exhibit symbiotic potential; in other words, those symbionts codify and amplify the PFP. For simplicity, our quantitative simulations using Lotka-Volterra equations do not explicitly model the recruitment of new symbionts. But the simulations are compatible with the life cycle of symbionts. And the equations are such that they can be easily extended to modelling also the selection of new symbionts. However, the parasite population is kept stable – around an equilibrium – by the Lotka-Volterra predator-prey relationship between prokaryotes and their viruses. Step 7: therefore, parasites qualify as catalysts, since i) the specific process is taking place in their absence, ii) if they are added the process is amplified, and iii) their population size is kept stable through-out and after the process. Step 7 implies that parasites are ubiquitous, and therefore, a host is a host-parasite, the latter being parasitized (by phages). Prokaryote-phage pairs are mutually stabilized within the host-parasite unit, as shown by the Lotka-Volterra equations. Steps 8-10: the two aforementioned energies act as counter force to parasitic virulence (step 8); for appropriate process stabilization the second, stored energy is required (step 9); finally and importantly (step 10), parasites do require to codify all relevant energy transformations, i.e. *E*_2_ → *E*_1_, *E*_2_ → *E*_1_ and *E*_2_ → *E*_1_. Collectively, we established a ten-step model comprising a life cycle of three host-parasite units that build a catalytic metabolism. This metabolism captures, channels and transforms energy used by the host to resist the fitness erosion caused by mutation and genetic drift and to adapt to environmental stochasticity (*21, 31, 33*).

## Discussion

Our model capitalizes on the growing body of knowledge demonstrating the ubiquitous nature of genetic parasites and their inextricable ties to genome replication, and therefore to all life forms, including LUCA and pre-LUCA organisms. Furthermore, host-parasite co-evolution is considered a major driving force of biological innovation and diversity.

This work firstly, confirms and extends previous findings by showing that parasites stabilize their host, modelling the host implicitly; secondly, it directly links – to the best of our knowledge for the first time – the central metabolic parameter, energy, with parasite abundance and host fitness. Thirdly, we provide a problem solving paradigm how adaptive processes involving host-parasites linked to their energy resources and fitness potentially materialize and evolve, an additional innovation. Future work will expand this model to address additional complexities including the effects exerted by direct and indirect anti-viral mechanisms such as virophages and antiviral responses, and their relevance for virulence and viral reservoirs.

## Methods

### Fitness

Let *P* be a quasispecies with *N* subpopulations *P* = (*P*_1_, *P*_2_, …, *P_N_*)^T^. *m* = *m_i_* denotes the (Malthusian) fitness: 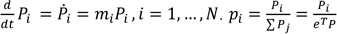 denotes the frequencies of the subpopulations *P_i_* within quasispecies. 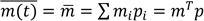 is the average fitness and 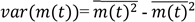 is the fitness variance of the quasispecies at time *t*.

### Fitness of parasites

The term *a*_1_ > 0 is the fitness of prokaryote *X* within its host habitat in the absence of phage *Y*. *a*_2_ > 0 is the fitness-decrease of the prokaryote *X* infected by phage *Y*. The term *b*_1_ < 0 is the negative fitness of phage *Y* in the absence of nutrition, i.e. prokaryote *X*, within the host habitat. −*b*_2_ > 0 is the fitness-increase of phage *Y* after having infected *X*.

### Virulence

Virulence is defined in its broadest sense, i.e. the cost to the host due to infection, which translates into the reduction in host fitness due to the infection (*34*). A parasite is said to be virulent in its host if it is actively reproducing. We assign values between 0 and 1 to the term virulence, virulence *v* = 1 if the parasite is maximally active and reproducing, and virulence *v* = 0 if the parasite is inactive and silenced.

### Energy

At least two energy forms exist to fight against the fitness decay of a quasispecies, *E*_1_is a form of immediately available energy (operating power, nutrition) that the quasispecies can use to directly fight erosion, e.g. induced by mutation genetic drift (*21*) environmental changes, and for reproduction. *E*_2_ is a form of stored, excess-energy *E*_1_, which in addition has an isolating effect towards the quasispecies’ inner and outer environment. This isolating effect partially neutralizes fitness erosion. Fitness increases/decreases as the level of both types of energy increases/decreases.

### One energy, one host-parasite unit homeostasis model

Virulence *v* of prokaryote *X* and its phage *Y*, in dependence of the energy level *E*:

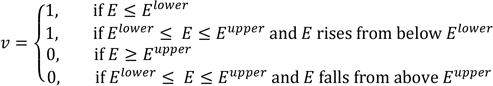

Lotka-Volterra equations of prokaryote *X* and phage *Y* under virulence 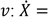

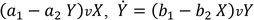, with equilibria 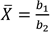 and 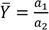, where *a*_1_ > 0, *a*_2_ > 0, *b*_2_ < 0, *b*_2_ < 0

Energy consumption and production differential equation: 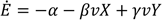, where

α > 0 is the rate of permanent energy consumption of the host-parasite unit,

β > 0 is the rate of energy consumption of parasite *X*, when virulent,

γ > 0 is the rate of catalytic energy production of phage *Y*, when virulent.

In this model we assume that from the prokaryote-phage pair (*X*, *Y*), only the phage *Y* catalyzes new energy *E*. Other possible realizations would be that both, *X* and *Y*, catalyze *E*, or only *X* catalyzes *E*.

As the energy level decreases below a lower threshold (*E* < *E*^*lower*^) the host-parasite unit gains full virulence (*v* = 1), indicating an evolutionary problem. The system now enters into the resolution search process (RSP) mode. The underling formalism related to evolutionary problem solving (*35*) is described below. If a sufficiently large population of symbiotic parasites is found fast enough, the system can recover. Fast enough means before the operating power *E* has reached zero. System recovery means that at least one element of the host-parasite unit catalyzes enough new operating power *E*, such that the energy level can rise again, using the available components of the host habitat and its environment. As soon as the energy level attains an upper limit (*E* ≥ *E^upper^*), the host turns fitter and regains the ability to tame and silence its parasites (*v* = 0). At this point, inactive symbiotic parasites cease to catalyze new operating energy. Therefore, the energy level declines below the lower level (*E* < *E*^*lower*^). The system faces again a problematic evolutionary situation, and parasites regain virulence. New energy will be catalyzed, and so on.

### Evolutionary problem solving

A quasispecies faces an evolutionary problem (i.e. mutation, genetic drift, environmental change) when the range of Malthusian fitness values for all its subpopulations *P_i_* is negative (*m*_*i*_ < 0, *i* = 1, …, *N*). For evolutionary problems we use a modified formalism based on classical problem solving in mathematics (*35*), namely i) a problem measurement phase (“understanding the problem”), ii) a problem resolution phase (“devising and carrying out a plan”), and iii) a fine-tuning phase (“looking back”).

Applied to our model, evolutionary problem solving can be contextualized as follows. Fitness decay, e.g. due to mutation and genetic drift, not only contributes to intensify the evolutionary problem, but also contributes to its resolution. We call this process RSP. Commonly, the positive effect exerted by fitness erosion via increased variance is rather weak, compared to its negative, destructive effect. Parasites typically amplify the RSP process by accentuating destruction and increasing the fitness variance. If this amplification is strong enough – such that the right tail of the quasispecies’ fitness distribution shifts well into the positive territory – then the negative, destructive effect becomes irrelevant.

If parasites survive and preserve their ability to amplify the RSP process, they then also codify it, i.e. they are stable information carriers of the RSP amplification process. Provided that the resolution research process RSP – via broadening of fitness variance – is amplified strongly enough, moving some host members of the quasispecies into the positive fitness range, implies that the following must have happened. Some of the parasites became symbiotic and therefore produce a positive fitness increase that is exerted both on themselves and on their hosts. Let us call this process problem fixing process (PFP). This subgroup of now symbiotic parasites carries the essential information of the resolution and PFP process, specifically reflecting the fitness problem the quasispecies has been facing.

### Two energies, three host-parasite units homeostasis model

The following three processes have to be codified and catalyzed in the quasispecies host-parasite unit:

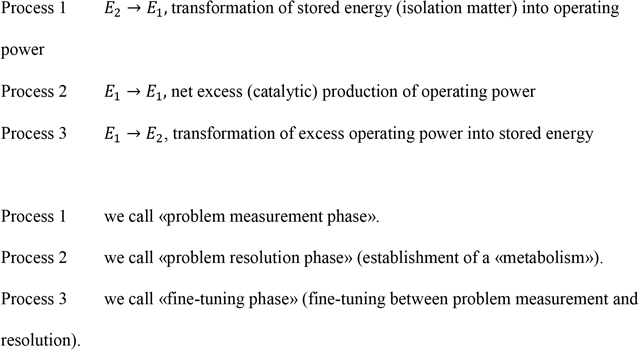

For a quasispecies system are given two energy levels (in each host) (*E*_1_, *E*_2_) and three pro-karyote-phage pairs (*X*_1_, *Y*_1_), (*X*_2_, *Y*_2_) (*X*_1_, *Y*_2_). In our model realization, the virulence of the pair (*X*_1_, *Y*_1_) is regulated by *E*_1_ (virulence 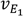) and the virulence of the pairs (*X*_2_, *Y*_2_), (*X*_3_, *Y*_3_) is regulated by *E*_2_ (virulence 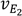).

Virulence 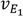 of parasite *X*_1_ and its phage *Y*_1_, in dependence on energy level *E*_1_:

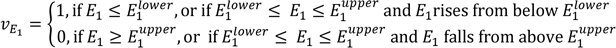

Virulence 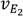 of prokaryote-phage pairs (*X*_2_, *Y*_2_), (*X*_3_, *Y*_3_), in dependence of energy level *E*_2_:

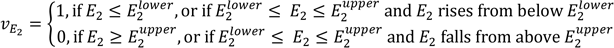

Lotka-Volterra equations (under virulence) of the prokaryote-virus pairs (*X*_1_, *Y*_1_), (*X*_2_, *Y*_2_) (*X*_3_, *Y*_3_):

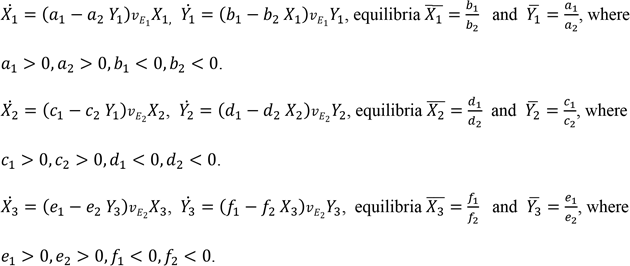

Energy consumption and production differential equations:

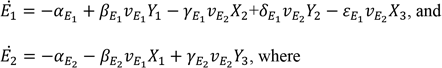

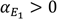 is the rate of permanent energy *E*_1_ consumption of the (host-parasite) system.

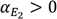 is the rate of permanent decay of natural isolation material in the (host-parasite) system.

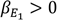 is the rate of energy *E*_1_ production of phage *Y*_1_, when virulent.

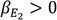 is the rate of energy *E*_2_ consumption of parasite *X*_1_, when virulent.

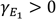 is the rate of energy *E*_1_ consumption of parasite *X*_2_, when virulent.

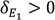 is the rate of energy *E*_1_ production of phage *Y*_2_, when virulent.

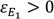 is the rate of energy *E*_1_ consumption of parasite *X*_3_, when virulent.

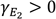 is the rate of energy *E*_2_ production of phage *Y*_3_, when virulent.

Thus, in this model the parasite-phage-pairs are information carriers for the following processes:

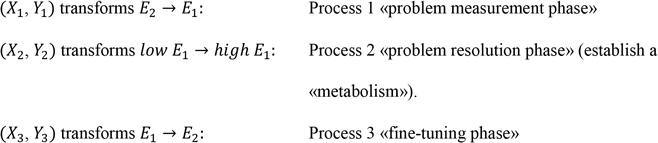

The first pair (*X*_1_, *Y*_1_) catalyzes Process 1, *E*_2_ → *E*_1_, transforming stored energy into operating power during the problem measurement phase. Virulence of (*X*_1_, *Y*_1_) is regulated by the operating energy *E*_1_. As the operating energy drops below a certain threshold 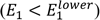, the system enters into a first stage of the problematic phase. The (*X*_1_, *Y*_1_) unit acquires virulence and transforms stored energy into operating power *E*_2_ → *E*_1_. This buys time for the RSP. If stored energy *E*_2_ decreases below a certain level 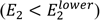, all *E*_2_– sensitive parasites acquire virulence. In our model, the host-parasite units (*X*_2_, *Y*_2_) and (*X*_3_, *Y*_3_) are *E*_2_– sensitive. The (*X*_2_, *Y*_2_) pair catalyzes Process 2, *E*_1_ → *E*_1_, leading to a net excess production of operating power in the problem resolution phase. And pair (*X*_3_, *Y*_3_) catalyzes Process 3, *E*_1_ → *E*_2_, the transformation of excess operating power into stored energy in the fine-tuning phase. The selection of an appropriate fine-tuning process is key, because it regulates the balance between isolation and influx of the components that (*X*_2_, *Y*_2_) depend upon to catalyze *E*_1_.

## Acknowledgments

We warmly thank Dr. Séverine Bontron and Prof. Joseph Curran, University of Geneva, for critically reading the manuscript, and we are indebted to P. Norbert Widmer OSB† for his inspirational and fundamental insights.

## Author Contributions

The manuscript was designed and written in a joined effort. Bernard Conrad contributed the biological background and modeling, Magnus Pirovino the mathematical modeling.

---

**Suppl. Fig. 1.**
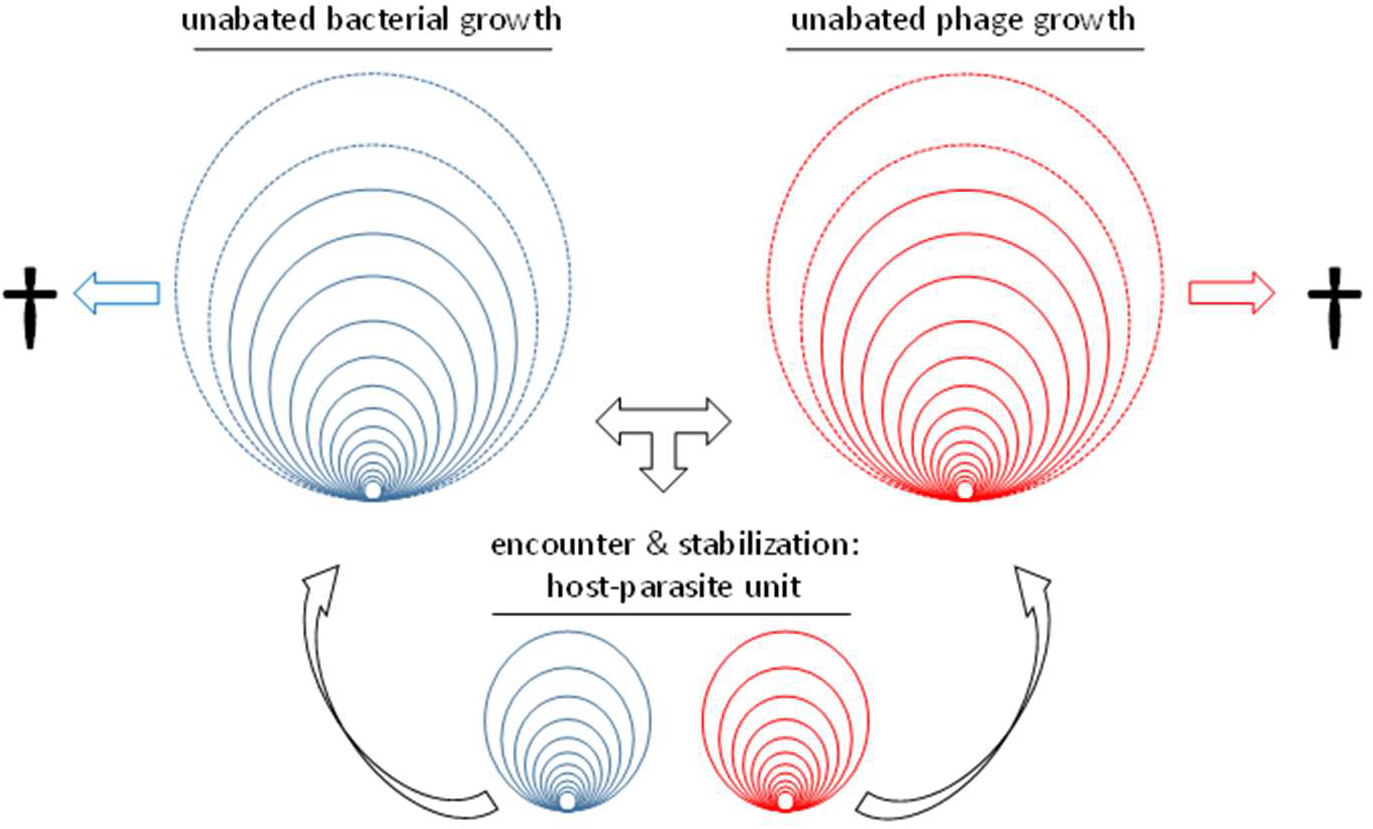
Host-parasite unit: parasite growth is unlimited (bacteria, phage), as long as the habitat nourishes it. Once the host loses the ability to nourish the parasites, they are condemned to perish. The association of a dedicated phage with a given parasite (bacteria) is a very effective and robust taming strategy for a parasitized host (both populations fluctuate around their equilibria).

**Suppl. Fig. 2.**
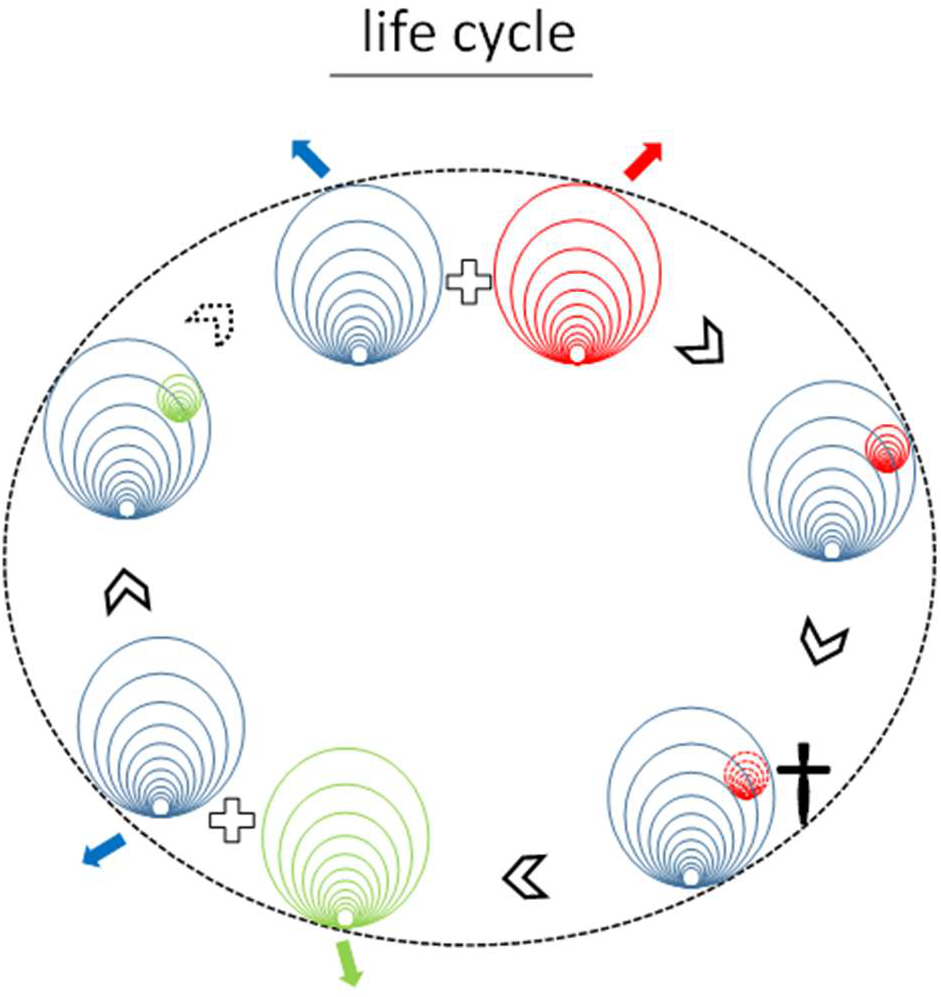
Catalytic host-parasite life cycle: an initially stabilized host-parasite unit recruits symbionts (red) that catalyze energy transformation. Degenerate endosymbionts are replaced by fresh symbionts (green) that catalyze the next transformative round. A trio of such host-parasite units and two energies are required for adequate system stabilization and to build a catalytic life cycle.

